# Physical Interactions Driving the Activation/Inhibition of Calcium/Calmodulin Dependent Protein Kinase II

**DOI:** 10.1101/2020.03.27.012393

**Authors:** Eliana K. Asciutto, Sergio Pantano, Ignacio J. General

**Affiliations:** School of Science and Technology, Universidad Nacional de San Martin, and CONICET, 25 de Mayo y Francia, San Martín, 1650 Buenos Aires, Argentina; Biomolecular Simulations Group, Institut Pasteur de Montevideo, Mataojo 2020, CP 11400 Montevideo, Uruguay

**Keywords:** electrostatics, phosphorylation, molecular dynamics, coarse-grained

## Abstract

CaMKII is a protein kinase whose function is regulated by the binding of the Calcium/Calmodulin complex (Ca^2+^/CaM). It is a major player in the Long Term Potentiation process where it acts as a molecular switch, oscillating between inhibited and active conformations. The mechanism for the switching is thought to be initiated by Ca^2+^/CaM binding, which allows the trans-phosphorylation of a subunit of CaMKII by a neighboring kinase, leading to the active state of the system. A combination of all-atom and coarse-grained MD simulations with free energy calculations, led us to reveal an interplay of electrostatic forces exerted by Ca^2+^/CaM on CaMKII, which initiate the activation process. The highly electrically charged Ca^2+^/CaM neutralizes basic regions in the linker domain of CaMKII, facilitating its opening and consequent activation. The emerging picture of CaMKII’s behavior highlights the preponderance of electrostatic interactions, which are modulated by the presence of Ca^2+^/CaM and the phosphorylation of key sites.

## 1. Introduction

Calcium/calmodulin (Ca^2+^/CaM) dependent protein kinase II, or CaMKII, is a serine/threonine specific kinase, important in many different biological functions, such as calcium regulation [1], signal transduction in epithelia [2], the cell cycle [3], T-cell activation [4], and turnover of focal adhesions and cell motility [5]. A particular aspect of CaMKII, focus of many recent articles, is its function in the process of long-term potentiation (LTP), i.e., the enhancement of synapses related to memory formation [6] [7] [8].

The protein is arranged as a homo-multimer, with 8 to 14 subunits (SU), where each of them is composed of a C-terminal hub domain, an N-terminal kinase domain, and a linker domain connecting them. The kinase domain is subdivided in a C- and an N-terminal lobes, whose intersection forms a hinge, constituting the catalytic cleft. These SUs get organized in two rings, stacked one on top the other, as shown in Figure 1 for the dodecameric case. Notice that, for practical purposes, we are defining the linker as containing the regulatory segment, and not as previously done elsewhere[9]. The hubs constitute the stable core of the system, and the kinases, located on the outer side of the multimer and attached to the center through the flexible linkers, have more freedom in their motion. The SUs are usually hypothesized[9] as being able to interchange between two conformations: one inactive or autoinhibited, in which the kinase lies against the hub; the other, active, when the kinase domain separates from the hub and moves freely (but still connected through the linker) in the neighboring volume. It is this latter state the one associated with the high kinase activity of the domain, which can phosphorylate other molecules, via the interaction of its catalytic site.

**Figure 1:**
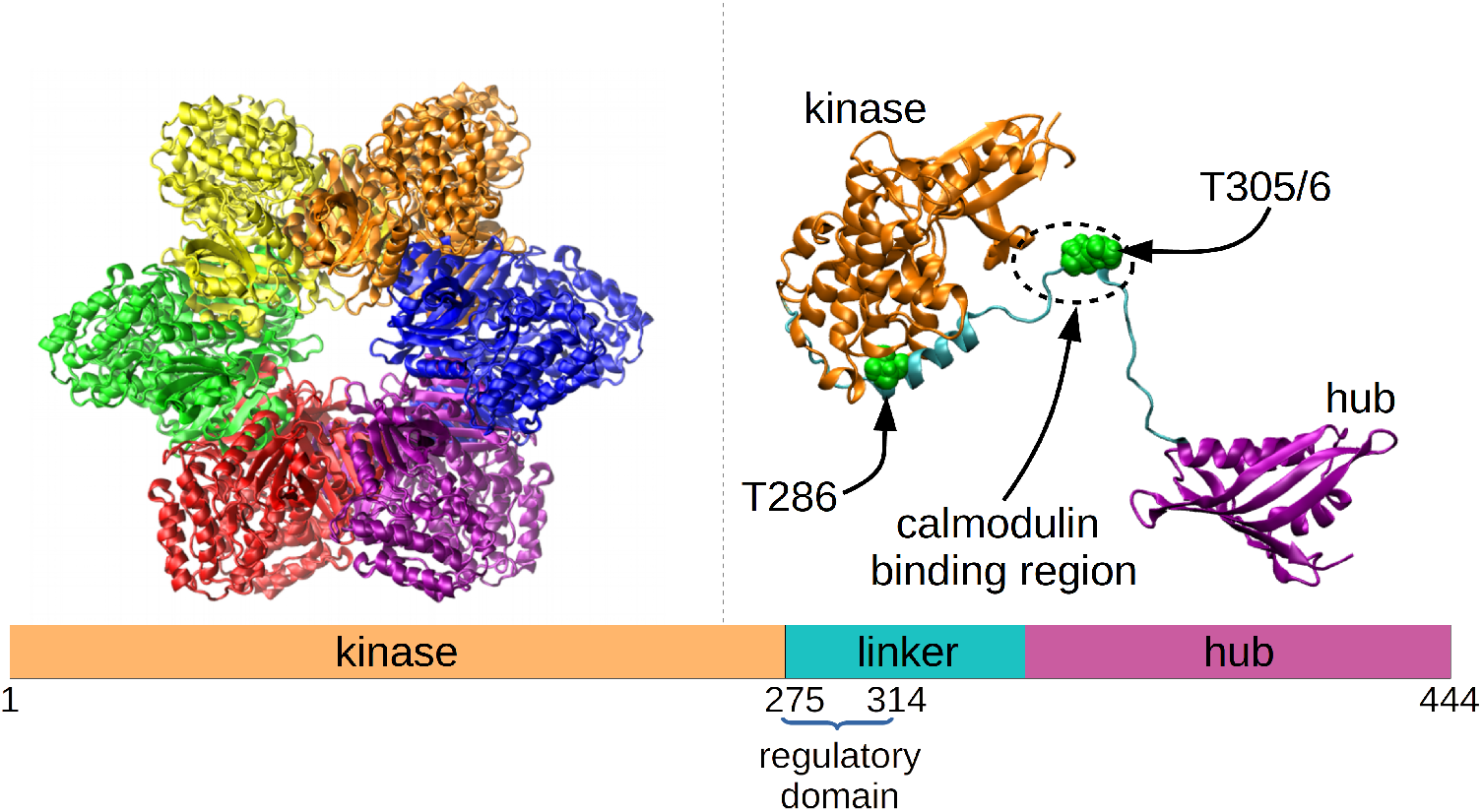
Modeled CaMKII dodecamer. It is a stack of two rings, each with six SUs, shown in different colors (one ring is behind the other, so they are not clearly distinguishable)(Left panel). Representation of one of the SUs, in a modelled open conformation, showing the hub and kinase domains connected via a flexible linker, and highlighting T286, T305 and T306–phosphorylation sites known to be critical for the protein’s function. At the bottom, there is a schematic view of the three subdomains (Right panel).

CaMKII belongs to a family of related isoforms, *α, β, γ* and *δ*, encoded by different genes[10]. The linker domain is subject to alternative splicing, resulting in many variants [11], all of them sharing a large sequence identity. Their most important difference is the length of the linker, which affects the frequency response of CaMKII to Ca^2+^/CaM pulses [12]. In this work, we study the predominant form of CaMKII found in the brain[13, 14], the α isoform, with a *β*7 linker connecting the C and N terminals. This linker has been shown to have a reduced kinase activity [15].

The LTP process is driven by the persistence of the kinase active state [16, 17]. The currently accepted picture describing the activation process begins with a SU whose kinase is docked onto the hub, in a closed inactive form. An incoming flux of Ca^2+^/CaM results in one of these molecules binding to the linker around residues T305-T306 [18], causing the displacement of the inhibitory regulatory segment and, in particular, releasing T286 from its occluded position against the kinase domain. Now, a more exposed T286 may become phosphorylated, inhibiting re-binding of the segment to the kinase domain and making the SU stay in an open, persistent, conformation. Once phosphorylated, not even unbinding of Ca^2+^/CaM can lead the system back to the original inactive state. In this way, Ca^2+^/CaM is needed to promote the activation, but it is not needed once T286 becomes phosphorylated. The kinase, away from the static hub, can move freely, interacting with neighboring SUs, and eventually phosphorylating them (trans-phosphorylation). Rebinding of Ca^2+^/CaM to T305-T306 is prevented by the phosphorylation of either residue[19, 20, 21, 22].

In a previous work [23], we found that each domain of the SUs of CaMKII contains a charged patch of residues: 1) Kinase (negative patch): E99, D100, E105, E109 and D111; 2) Linker (positive patch): K291, K292, R296, R297, K298 and K300; 3) Hub (negative patch): E325, E329, D335, E337, K341, D344, E360 and D363. These patches also have a high aminoacid sequence conservation, suggesting a strong structural and/or dynamical relevance. The role of phosphorylation–as put forward in that work–is to change the charge relation between the patches, turning on and off their interactions, and switching between attractive (non-phosphorylated or inhibited) and non-attractive (phosphorylated or active) states. More specifically, the natural distribution of these patches is hub-linker-kinase (Figure 1) or, in terms of their charges, negative-positive-negative. The positive patch in the center is fundamental in allowing the hub and kinase domains to stay close to each other, despite having like charges, as shown in the mentioned work. Without the positive linker patch, hub and kinase would repel each other. Consequently, binding of a strong negatively charged molecule (Ca^2+^/CaM) in the linker region, or phosphorylation of linker residues (T286, T305 and/or T306) reduces the linker’s positive charge, thus neutralizing the attractive state of the system, and favoring the hub-kinase separation.

In this work, we used multiscale Molecular Dynamics (MD) simulations, both at the all-atom (AA) and residue-based coarse-grained (CG) levels. The first case produces more accurate results, but precisely due to this, it is computationally costly. On the other hand, CG models are less accurate but allow much longer time spans of the simulations.

Here, combining both approaches in order to extract robust results from a local and a global perspective, we investigate the effect of different variants of the system (wild-type, phosphorylated, with ions, with Ca^2+^/CaM) on the initial steps of opening of the SUs and their later activation.

## 2. Methods

### 2.1. All-atom model of a CaMKII monomer (no CaM)

The crystal structure with pdb code 3SOA[9] describes human CaMKII in its *α* isoform, with a *β*7 linker, in a closed or auto-inhibited conformation. We used this structure to form the closed SUs for all our MD simulations.

With the objective of studying the influence of electrostatic forces in the process of SU opening, we built three AA models: the first one, the control system, in a solution of water (with two Na^+^ counter-ions); the second one in the same solution, but reducing the atomic charges of the charged residues in the positive patch of the linker (K291, K292, R296, R297, K298 and K300), so that they became neutral (this is an ideal case where we can test the relevance of the linker’s charge by going to the limiting case where it loses all its positive charge); and the third system in a 0.15M solution of 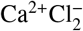. The three systems were electrically neutral. We should make clear that the biologically relevant system to study is not CaMKII with ions and with a charged or uncharged linker, but CaMKII with Ca^2+^/CaM, since this is the molecule known to regulate CaMKII’s activation. Such a simulation would require making very large systems of CaMKII and unbound Ca^2+^/CaM, that would take very long to run in an AA simulation. As an approximation to this, and motivated by our previous finding [23] that electrostatics is the driving force in CaMKII’s dynamics, we chose to study the interactions of CaMKII in a system where we emulate the interaction of a negatively charged unit (Ca^2+^/CaM) that tends to neutralize the positive charges in the linker, with a system where the linker is already neutral (our second system above). And in order to further probe the importance of electrostatic interactions, we also prepared the third system where positive ions could interact with the negative patches in hub and kinase (although there is no evidence that CaMKII is exposed to free Ca^2+^ ions, we believe this is a good way to test the relevance of these interactions).

Ions were modelled with the Joung/Cheatham [24] (Na^+^ and Cl^−^) and the Li/Merz [25] (Ca^2+^) parameters for TIP3P water. The simulation boxes were made in octahedral shapes, and such that the minimum distance from any residue of CaMKII to the wall of the box was, at least, 10 Å. Long-range electrostatics were taken into account using periodic boundary conditions, via the Particle Mesh Ewald algorithm [26], with a cut-off of the sums in direct space of 10 Å.

The protocol for driving the AA monomer model to a stable conformation consisted of several cycles of minimization and equilibration. The simulations in this work were performed using the AMBER16 software package[27], with all of them using the GPU version of the PMEMD program. We employed the ff14SB[28] force field. The system was kept at a temperature of 298 K, using Langevin dynamics with a collision frequency of 2 ps^−1^, and a pressure of 1 atm, via a weak-coupling Berendsen barostat, with a relaxation time of 2 ps. The SHAKE algorithm was adopted, allowing the use of a 2 fs time step. The protocol followed for minimization and equilibration was: 1) 1,000 cycles of minimization, using AMBER’s XMIN method, followed by 5,000 cycles using steepest descent, and another 5,000 steps using conjugate gradient; 2) 1 ns of heating, to 298 K, followed by another ns at constant temperature and pressure (1 atm), and finally 50 ns with constant temperature and volume.

The simulations of this section were performed using harmonic restraints, in order to drive the system to an open conformation (more details in next section). Several spring constants (*k* =0.01, 0.05, 0.1 and 0.5 kcal/(mol·Å)) were used, finding *k* = 0.1 kcal/(mol·Å) to be the most sensitive, and thus using it for the production runs: nine 500 ns simulations were performed for each environment, water, neutral linker and positive ions, for a total of 13.5 *μ*s.

### 2.2. All-atom model of a CaMKII monomer with two CaM molecules

Our goal in relation to this system was to drive an initially closed SU towards an open conformation, and analyze the binding of Ca^2+^/CaM on the open SU. Therefore, we took the previously described wild-type monomer, this time in a 0.15 M solution of Cl^−^Na^+^, and added two Ca^2+^/CaM molecules a few angstroms away from CaMKII (taken from the 2WEL pdb structure[29], which contains an open SU with bound-Ca^2+^CaM), one on each side of the grooves formed between the hub and kinase domains, and such that there was no contact between them. After applying the same equilibration protocol mentioned above, the SU was forced to open with a harmonic force with constant *k* = 0.1 kcal/(mol·Å), for 10 ns. Then, five independent 250 ns production runs, without restraints, were performed for a total of 1.25 *μ*s.

### 2.3. Coarse-grained model of a CaMKII monomer with unbound CaM

The models described next are CG systems, parametrized according to the SIRAH2.0 force-field for proteins [30]. SIRAH uses three beads per aminoacid backbone, and between zero and five beads for side-chains. It’s associated with an explicit water model, WT4, consisting of four beads per eleven water molecules, and it also contains parametrizations for several ions, including *Ca*^2+^. This force-field has been successfully applied to the study of diverse Calcium related systems[30, 31, 32].

We built a monomeric system, with CaM, as follows: starting with the 3SOA structure, we performed a short atomistic simulation where harmonic restraints forced the separation between the center-of-mass (CoM) of hub and kinase, in order to increase it to a value of about 100 Å. Next, a Ca^2+^/CaM molecule (same as used in section 2.2) was similarly opened–separating the two EF lobes (each lobe contains two pairs of EF hands) to a CoM distance of about 50 Å–and placed next to residues T305 and T306 of the monomer; this is the region where Ca^2+^/CaM has been observed to bind ([19, 33]). This AA system was later minimized and equilibrated for 12 ns, following the same protocols described previously; this was done using soft restraints to the coordinates of the just opened system, so that it could relax without leaving the initial conformation with both molecules opened, and Ca^2+^/CaM close to its binding region. With this relaxed AA system, we then coarse-grained it using SIRAH Tools[34], and ran another round of minimization/equilibration. The overall equilibration time was above 1 *μ*s, starting in the NPT (50 ns) and switching to the NVT ensembles. And finally, a 41 *μ*s production run was performed, always keeping restraints in the coordinates of the hub, in order to maintain it in position, imitating the restrictions of the hub’s motion when in presence of neighboring SUs. The temperature and pressure parameters used in the CG simulations were the same as those in AA runs.The time-step was set to 20 fs.

### 2.4. Coarse-grained model of CaMKII (no CaM)

Using the crystallographic symmetry matrix contained in the description of the 3SOA pdb structure, we built three dodecameric CaMKII models. One was the wild-type (WT); the second emulated a phosphorylation of T286 (PT286) in each of its SUs, by doing a double phospho-mimetic substitution, T286E and V287D. And the third one had the charges of all linker domains manipulated, in order to electrically neutralize them (NE), as was done for one of the AA systems previously described (see section 2.1). The protocol used to equilibrate each of these systems was the same described above, for the monomer. Afterwards, a production run of 38 *μ*s was performed for each case.

The final conformations of those runs were also used as a starting point for a free-energy calculation of the opening process. Using the Umbrella Sampling method, the system was driven to an open conformation following a reaction coordinate defined by the CoM separation between the kinase of a given SU and the rest of the system (hub and all other SUs). Its value started in the range 62-72 Å (range of values for the different systems, as observed in the CG simulations; see next section) and was increased to 140 Å (corresponding to SUs where their kinases are totally removed from the central hubs), by constraining the coordinate with harmonic potentials to move within windows centered at different consecutive values along the initial and final separations. Those windows were taken every 1 Å, and the magnitude of the harmonic constant used was 10 kcal/mol·Å^2^. The simulation time in each window was 40 ns, giving a total time of almost 3 *μ*s per umbrella sampling run. We performed two runs for each of the three systems.

## 3. Results

### 3.1. Neutralization of electrostatic interactions favors subunit opening

Why is it that an initially inactive (closed) subunit of CaMKII opens, thus allowing the phosphorylation of T286? Ca^2+^/CaM is thought to be the causing factor of this behavior, but what are the molecular mechanisms involved? Sutoo and Akiyama [35] studied the clinical effects of administering 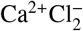 or 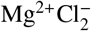 to sleeping mice, and noticed a strong difference in their response, suggesting that the Ca^2+^ cations have important effects in the brain, probably via interaction with CaM. Related, Zhang *et al* [36] found that the Ca^2+^/CaM complex binds with an overwhelmingly higher affinity to CaMKII than Ca^2+^-free CaM. Inspired by these ideas, we hypothesize that the ultimate explanation to CaMKII’s activation is the electrostatic interaction between the charged patches of the SUs, mentioned in Pullara *et al* [23] (negative patches in hub and kinase, and positive in linker), and the negative Ca^2+^/CaM molecules. In order to investigate this, we performed three sets of AA MD simulations of a CaMKII monomer, without CaM, in its closed form. These are the systems described in section 2.1: the control case, in aqueous solution; a study case in aqueous solution, but artificially neutralizing the positive residues in the linker to produce zero net charge (this case was devised to emulate the electrostatic consequence of Ca^2+^/CaMKII binding to the linker). Finally, a third system was prepared in presence of a 0.15M solution of 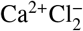 (with wild-type fully-charged linker). Since computationally reasonable times of hundreds of *ns* are not enough to show significant differences between the three sets, we decided to drive the systems to an open conformation, using harmonic constraints (springs) between the CoM of the hub and kinase domains. In Figure 2 we present the kinase-hub inter-domain distance for nine independent simulations of the three systems, all with the same harmonic forces pushing the system open, toward a target distance of 80 Å. In the left panel, blue and red lines were used for cases with the mutated neutral and the wild-type charged linkers, respectively. On the right side of the panel, we observe a histogram of the last 200 ns of simulations, when the systems were well stabilized. Equivalently, the right panel shows the results for the charged linker without Ca^2+^ ions (same run as in left panel), and the same linker with Ca^2+^ ions present in the solution (blue lines). The two panels show similar results, in the sense that the altered systems (neutral linker and presence of cations) are driven more easily to open conformations: the blue histograms are located above (larger separations) the red ones.

**Figure 2:**
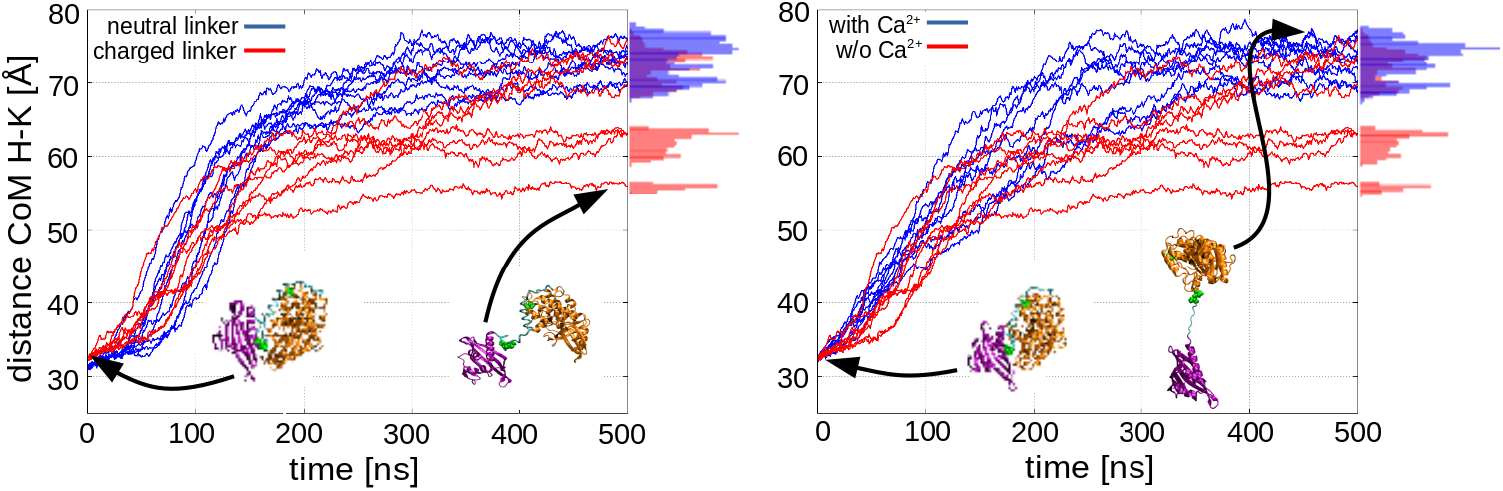
When pulled by a harmonic force, a monomer SU tends to have larger inter-domain separations when the linker is electrically neutral (blue curves, left panel), or there are Ca^2+^ ions in the solution (blue curves, right panel), as compared to a wild-type linker with no Ca^2+^ ions (both panels, red curves). Conformations during the trajectories show a more stretched linker in the cases represented with blue curves

In regard to the molecular mechanism driving this behavior, the explanation appears clear in terms of the charged patches interpretation of our previous study [23]: 1) A neutralized linker leaves two negative patches directly interacting, as it removes the intermediate positive patch, which acted as an attractor to both of them. 2) Added positive ions tend to interact with the negative residues in the hub and kinase patches, thus partially neutralizing them (MD trajectories show this; see left panel of Figure 3), and again, breaking the attractive electrostatic interaction between the three patches. Although the concentration of Ca^2+^ ions may not reflect physiological conditions, this result highlights the relevance of electrostatic interactions. In summary, both scenarios cause a partial neutralization of the attractive interaction between patches, facilitating the hub-kinase separation.

**Figure 3:**
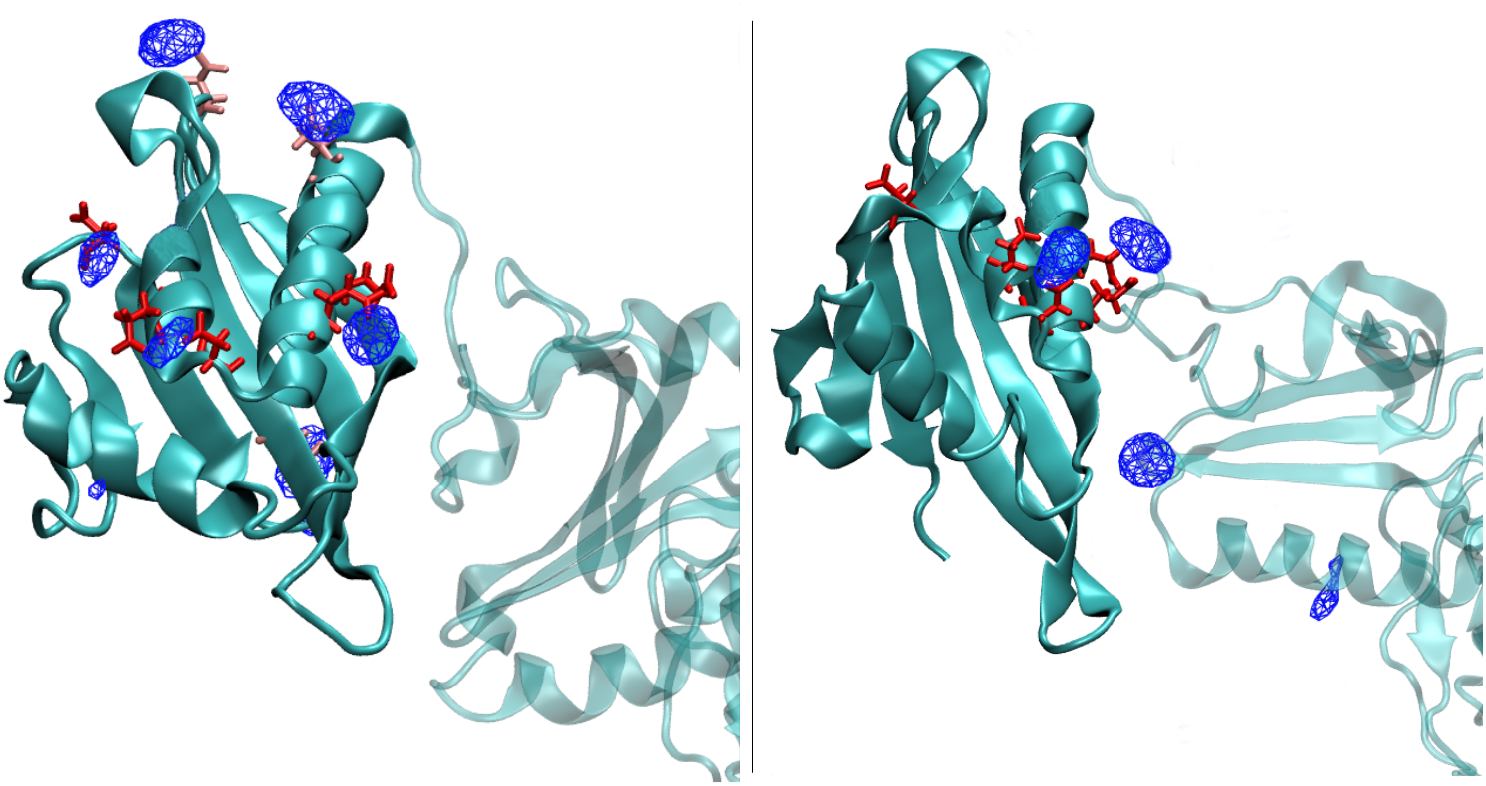
Cations tend to quickly bind to negative residues in the hub domain (solid cyan), weakening the attraction between charged patches in hub, linker and kinase domains (last two in translucent cyan). Blue mesh objects represent the regions where positive ions are found most frequently. Left panel: In the simulations without CaM, the cations strongly bind to residues E325, E329, D335 and E337 (in red licorice). They also tend to bind to residues E319 and D394 (in light red), although these two are not part of the hub’s negative patch. Right panel: the simulations with CaM (not shown) also show that cations tend to strongly bind to the negative patch of the hub.

The large separation between hub and kinase domains at the end of the opening simulations (500 ns left panel of Figure 2), about 75 Å, may seem too large for electrostatics to be significant. But a rough estimate for the strength of this force between the pairs of patches gives magnitudes of 0.1, 0.2 and 0.4 kcal/(mol·Å), for hub-kinase, linker-hub and linker-kinase, respectively, considering the screening effect of water without ions. These numbers are to be compared with the harmonic force of the restraint, around 0.5 kcal/(mol·Å), corresponding to a spring stretching of 5 Å (also for the end times of the opening simulations). We see, thus, that the two types of forces, harmonic and electrostatic, are comparable, and the natural attraction between patches is still relevant. On the other hand, when the linker residues are neutralized, the only remaining electrostatic interaction is the repulsion between hub and kinase; so it just adds to the harmonic restraint; and when Ca^2+^ ions are introduced, the screening effect gets highly increased (Coulomb forces decrease exponentially for large separations) and the harmonic force becomes the main driving interaction. These facts explain the difference between the blue and red curves in Figure 2.

It is also worth mentioning that these are out-of-equilibrium simulations, since they were done with an external force pushing the system open in a short time. But this is not a problem in terms of the conclusions we draw, which are based on the internal forces of the system, particularly on electrostatics. Nevertheless, in order to confirm our results, we repeated the simulations using a much softer thermostat, with a coupling constant of 0.2 *ps*^−1^, which is more appropriate for out-of-equilibrium simulations[37] (although temperature could have larger fluctuations). We obtained practically identical results to those in Figure 2, highlighting the robustness of our approach to capture the physical effects on the system.

Despite the satisfactory description obtained, the computational experiments described considered one single CaMKII chain. However, in the physiological context, kinases in a multimeric array may interact not only with their own hubs, but also with those from neighboring proteins, and their interactions may affect the way in which the system behaves globally. To investigate this further, we used the Umbrella Sampling method of free-energy calculation for the study of the opening of three CG dodecameric structures, all with initially closed SUs, but differing in their electrical charges; these are the systems described in section 2.4, WT, PT286 and NE. We calculated the free-energy profiles for the opening of a SU, starting from equilibrium conformations (CG structures after 38 *μ*s simulations, as described later in section 3.3). The left panel of Figure 4 shows the results, where we can observe that both PT286 and NE show a plateau starting around 100 Å, unlike WT which shows no sign of reaching any kind of stability along the process. The structure in the Figure shows a representative conformation of the system around the plateau, where the harmonically forced SU appears interacting with a neighboring kinase. We also performed a free energy calculation in the reverse direction, reducing the hub-kinase separation of the SU, starting from the last frame of the corresponding opening simulations. This is showed in the right panel of the Figure where, interestingly, the minima are not observed for separations of about 70 Å, as was the original case, but they appear at significantly larger values, above 100 Å for PT286 and NE. The reason for this shift in position is clear when one observes the actual MD trajectories: the open kinase cannot go back to its original location–docked to its corresponding hub–due to steric clashes with its neighbors and, instead, it stays slightly out of the ring, and tilted to a side, approaching a neighboring SU. The structure in the Figure (from rotated points of view) shows a conformation corresponding to the region with minimum free energy. Notice that even in this case, with domains crossed, WT shows a hub-kinase separation shorter that the rest, still expressing the strongest attraction due to its fully charged domains.

**Figure 4:**
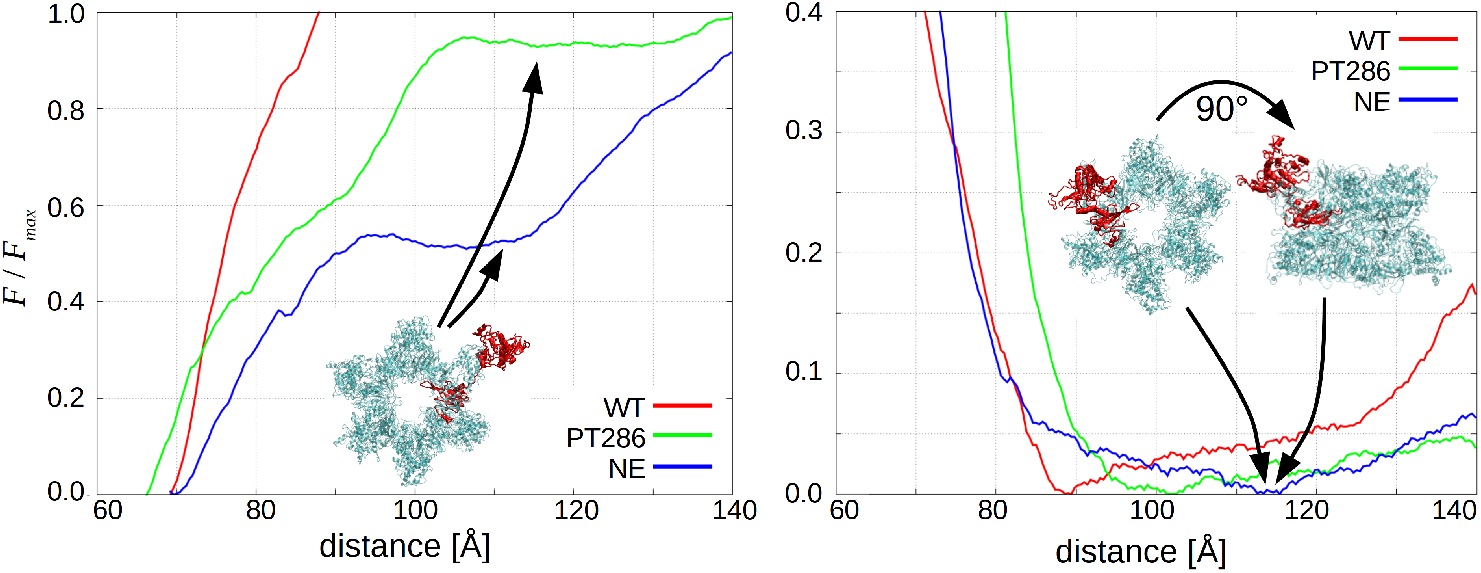
Free energy profiles of opening (left panel) and closing (right panel) of a SU for each simulated case (wild-type, phosphorylated, and neutralized linker), shown as a fraction of the maximum value of free energy, *F*_*max*_. Errors=±0.1. Also shown are representative structures of the PT286 and NE systems, showing the forced SU in red.

It is worth to note that these free-energy calculations are performed on a CG system. Since the loss of degrees of freedom–intrinsic to coarse-graining–produces a generally flatter energy landscape, the values in Figure 4 provide only a qualitative picture of the energetics of opening/closing in a biologically relevant ensemble. With this in mind, we note a general trend: although the three studied cases tend to stay in a closed conformation if they are in it already (left panel, Figure 4), NE is the easiest to be opened to a metastable state, requiring about 0.5 *F*_*max*_ (where *F*_*max*_ is the largest value shown in the plot for WT); it is followed by PT286, with 0.9 *F*_*max*_; and finally WT, which fails to reach such a state, increasing monotonically until the maximum distance. On the other hand, when the SUs are already opened (right panel, Figure 4), NE stabilizes with the largest hub-kinase separation, followed by PT286 and, finally, WT. The trend is the same in both directions, supporting what was previously suggested: the electrostatic interactions set up by the charged patches of CaMKII are responsible for keeping the SUs closed and, thus, the disruption of this equilibrium, via different mechanisms (phosphorylation, neutralization of the linker or addition of cations), enhances the probability of the system switching to an open conformation.

### 3.2. CaM binding to linker domain

We also investigated the binding of CaM to the CaMKII enzyme, starting from an AA wild-type monomer, in the closed conformation, this time adding two Ca^2+^/CaM molecules, not initially bound to CaMKII (described in section 2.2). As in a previous case, in the absence of CaM, we observed the cations quickly partitioning in the close neighborhood of the residues in the hub’s negatively charged patch, with binding times on the order of 10 ns (Figure 3, right panel). Again, this causes a partial neutralization of the attractive interaction between patches, favoring the hub-kinase separation.

The opening process, necessary for CaM to get to its binding region in CaMKII (see Figure 1), appears to be stochastic and can, thus, take a very long simulation time. Therefore, and as done in the previous section, we employed harmonic restraints in order to drive the system to the open conformation. A harmonic potential with spring constant of 1 kcal/(mol·Å) was applied between the CoM of the kinase and hub domains. The CoM separation grew from 33 to 90 Å in about 10 ns. Next, with the system in this open conformation, we performed a 250 ns MD, without constraints. We repeated this process five times, obtaining five independent runs.

In all performed simulations, the CaM molecules quickly attached around the linker region, sterically forbidding the re-docking of hub and kinase. The left panel of Figure 5 shows a significant number of hydrogen bonds formed between either of the CaM molecules and the kinase domain, underlining their strong interaction. With CaM in contact with the kinase and preventing the kinase-hub re-closure, the flexible linker is left to explore the volume between the two subdomains. When the region around T305/306 gets close to CaM, some hydrogen bonds are formed (right panel of Figure 5), and CaM stays bound to the linker, around those two residues. In particular, in one of the runs an average of six bonds were present at any time, indicating a rather strong linker-CaM interaction. The table in Figure 5 indicates the most active linker residues in forming hydrogen bonds with CaM. This is consistently repeated in all five simulations. Figure 6 portraits a representative conformation of the system when the two CaM molecules are forming hydrogen bonds with the region of the linker around T305/306. Another interesting observation is the activity of residues K21 and R415, located in the kinase and hub domains, respectively. These aminoacids were found to interact strongly with CaM for about 80% of the total time.

**Figure 5:**
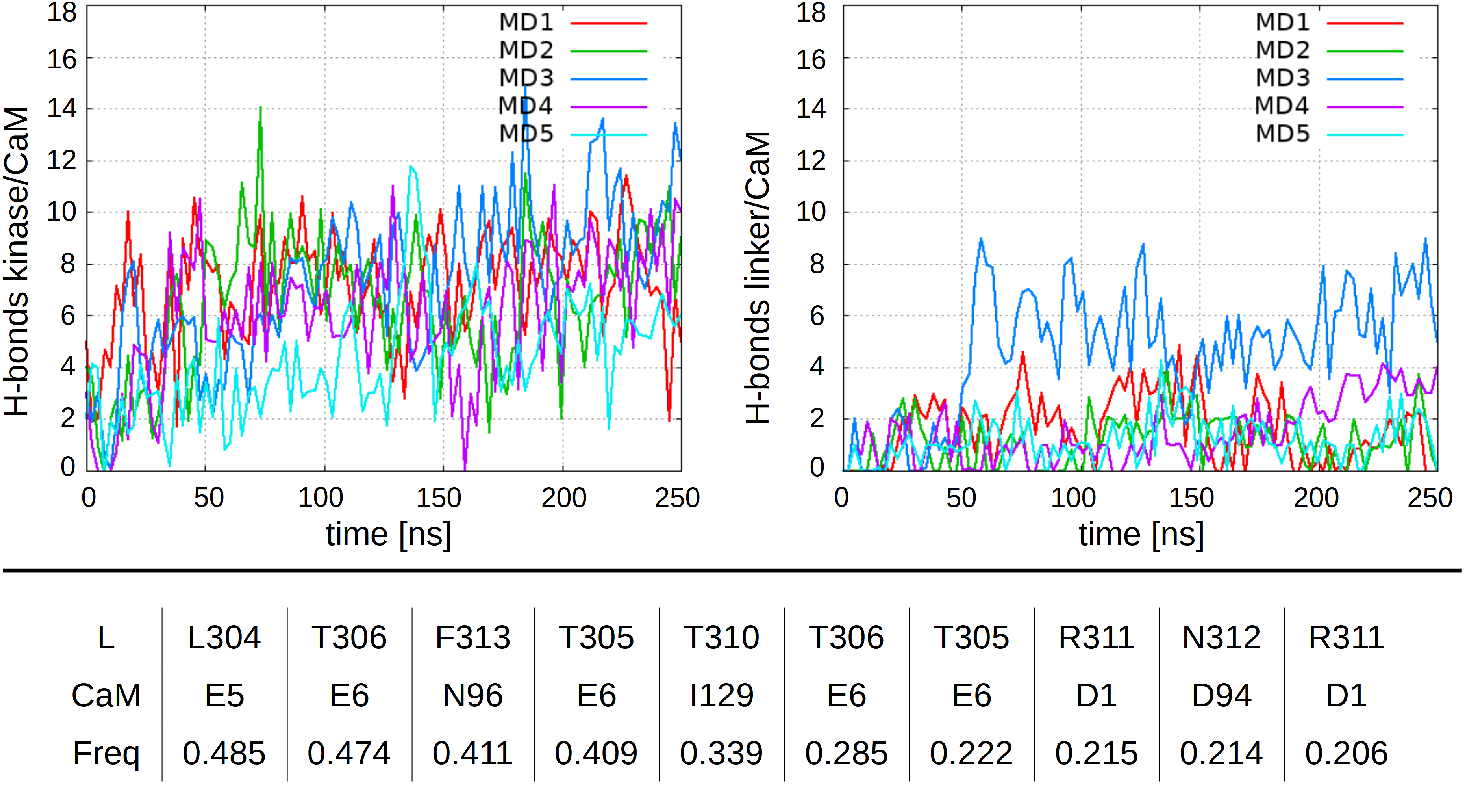
Top panel: Number of H-bonds formed between any Calmodulin molecule, and the linker or kinase domains, in each of the five simulations. Bottom panel: Linker-CaM residue pairs forming hydrogen bonds, and their frequency of occurrence along the trajectory (each column represents the contribution of a single CaM molecule in a single trajectory)

**Figure 6:**
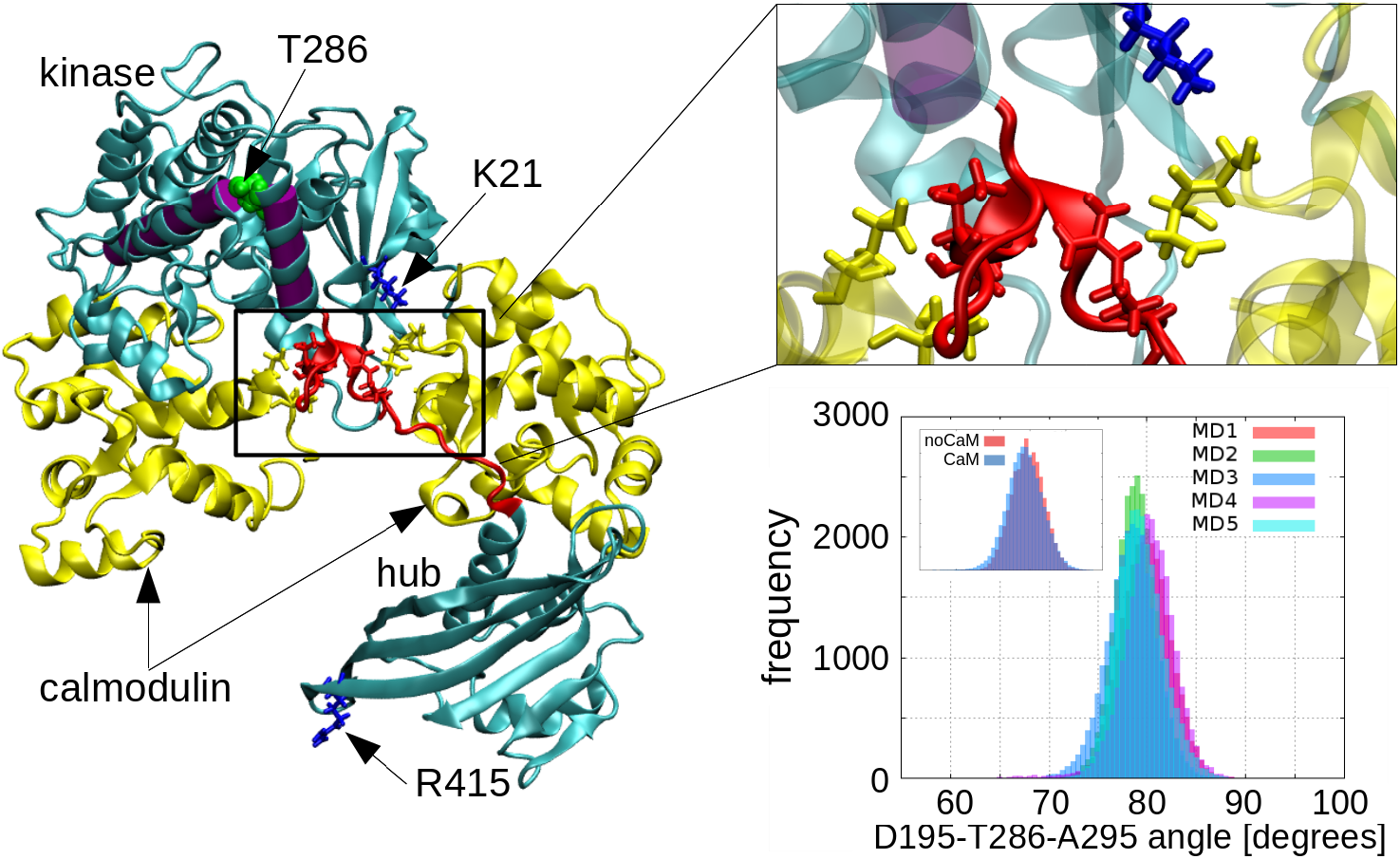
Kinase and hub domains in an open conformation, with two Calmodulin molecules (yellow) interacting with the linker (red), forbidding their re-closure. Residues K21 and R415, in blue licorice, show strong interactions with CaM (see text). The image on the top-right shows a close-up view of the residues forming hydrogen bonds, including aminoacids T305 and T306, in red licorice. The histogram on the bottom-right shows the variation of the D195-T286-A295 angle (angle between purple cylinders on the left) along the five MD trajectories

We should notice that the literature mentions that the mechanism of action of Ca^2+^/CaM to kick-start the activation process is to bind to the T305/306 region, hence preventing the kinase-hub re-closure. At the same time, due to its size, CaM pulls the regulatory segment out of its position docked to the kinase, relieving T286 from its enclosure against an internal mostly hydrophobic pocket–formed by L206, L207, V208, G209, Y210, E236, W237 and Y107– and becoming exposed to the exterior. In this way, a nearby kinase could interact with it and, eventually, phosphorylate it. In line with this, we measured a significant fluctuation of the inter-helix angle D195-T286-A295 (see angle formed by the purple helices H282-L299 and G191-V208, in Figure 6), which is indicative of a higher propensity of the regulatory segment to the opening. The histogram in Figure 6 shows this fluctuation–which is reproduced in all five runs, suggesting a robust sampling–spanning between 67° and 89°. The histogram corresponding to the simulation without CaM (inset in the figure) is very similar, with a slightly narrower distribution, with angles in the range 73° to 87°. Hence, we hypothesize the phosphorylation of T286 becomes possible not as a result of a variation of the inter-helix angle and consequent increase in exposure of T286, but as a result of a different mechanism related to the hub-kinase separation, to be explored below with CG simulations.

CaM failed to arrive at a fully bound conformation with CaMKII in the CG simulations described in section 2.3, although some notable facts could still be observed. One of the EF hands of CaM did show a very close and stable association with the linker domain of the enzyme. This hand, which had been put initially very close to its bound position (following an analogous problem in [30]), stayed strongly associated to the region of the linker around residues T305/306, although its conformation did change, within 1 *μ*s, to a stable final state with an RMSD 6 Å away from the initial structure. The other EF hand, on the contrary, did not find a stable position in relation to the first hand, and freely moved in the volume between hub and kinase, making contact with residues in both of them. In particular, several microsecond-long interactions were registered between CaM and residues around K21 in the kinase domain, and R415 in the hub domain (see Figure 6). Interestingly, these two residues were also observed in the AA simulations to have strong interactions with CaM, as noted above. This coincidence between both models (AA and CG) suggests that K21 and R415 could be important actors during the binding process of CaM.

### 3.3. K21 and R415 as promoters of CaM binding

The starting points of the CG simulations were the three dodecameric structures already mentioned in section 2.4: wild-type (WT), phosphorylated (PT286) and neutralized linker (NE) versions of CaMKII. After a 38 *μ*s simulation of each system, none reached a fully open conformation, although about half of the SUs in each case did significantly increase its hub-kinase CoM separation: WT showed the smallest variations, reaching peaks of about 37 Å for one of its SUs; PT286 and NE showed significantly larger separations, reaching peaks of 42 and 44 Å respectively. These maximum openings for each case are shown in the right panel of Figure 7, while its left panel displays representative conformations for the phosphorylated case.

**Figure 7:**
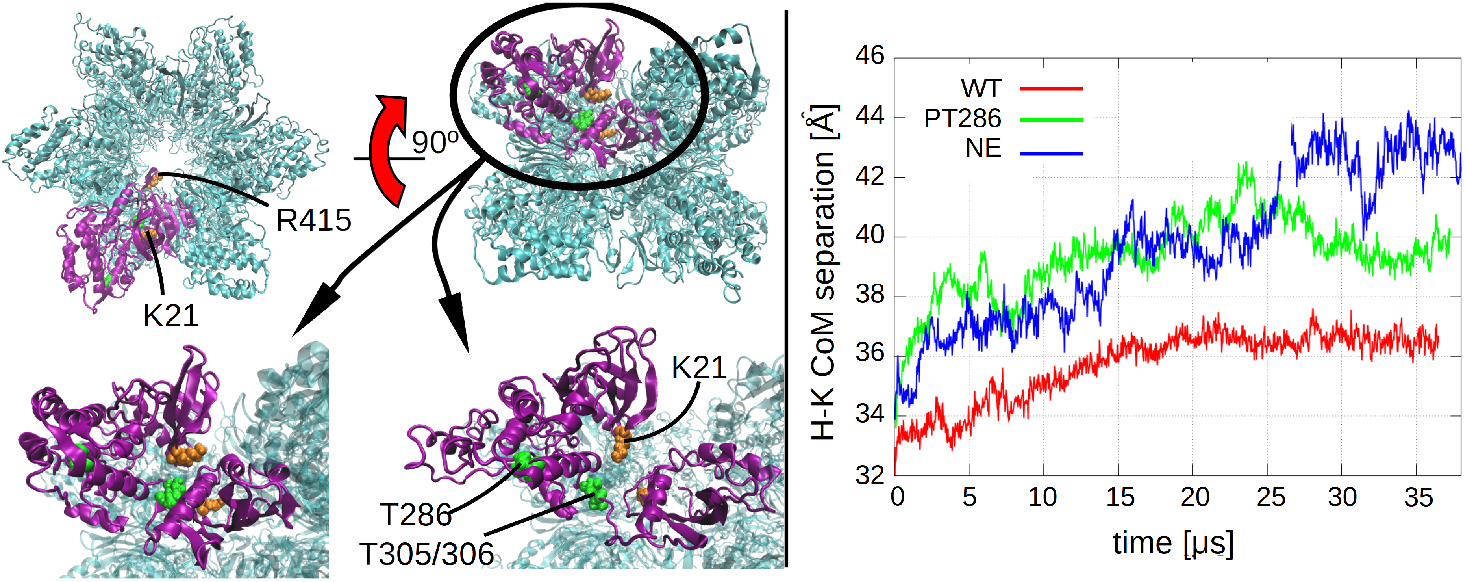
Left panel, top pictures: rotated views of the dodecamer, highlighting one of the SUs (purple), residues K21 and R415 (orange), and residues T286, T305/306 (green). Left panel, bottom pictures: zoomed-in views of the SU in the upper pictures, showing the initial (left) and final (right) conformations of one of the MD simulations. The hub-kinase separation increases, allowing K21 to become more exposed. T305/306 remains still partially buried. Right panel: the charge-altered cases, PT286 and NE, show the largest (center of mass) hub-kinase separation

On inspection of the conformations reached by the systems once their SUs become stable in terms of their inter-domain separations (last third of the simulation time), we find that K21, a neighbor of T305/T306, becomes more exposed than when in a closed form (compare bottom panels in Figure 7). But K21 was noted above as able to develop strong interactions with Ca^2+^/CaM. This leads us to hypothesize that K21 may act as a *bait* for Ca^2+^/CaM, bringing it closer to itself and its neighboring T305/306. Now being close, Ca^2+^/CaM may develop strong interactions with the latter residues (as in Figure 5).

A similar idea could be proposed for R415, since it was also noted previously as developing strong interactions with CaM, and it was also observed to become partially exposed during the CG runs. But since this residue’s distance to T305/306 is larger (minimum K21-T305/306 and R415-T305/306 separations ~ 10 and 20 Å, respectively), this option through R415 seems to be less effective (although a possibility not to be discarded). This mechanism appears to be more likely in the non wild-type cases, due to the maximal exposure of K21, but it may also be a possibility in the WT, since Figure 7 also shows an increased separation for this case.

## 4. Conclusion

In this study we performed AA and CG MD simulations, in order to study the mechanisms that kick-start the activation process of CaMKII, and guide its ensuing dynamics.

From the atomistic runs we observed that a disruption of the electrostatic interactions between hub-linker-kinase of a monomer, either via neutralization of linker or addition of cations, enhances the SU opening when pushed apart by a harmonic force. Despite the use of a biasing force introduced to speed up the process, this observation confirms the findings in Pullara *et al* [23]; namely: electrostatic interactions between hub-linker-kinase are fundamental in determining the structure and dynamics of the system. Therefore, any event that disrupts that electrostatic equilibrium (phosphorylation, ion binding, etc), changing the relation of charges between them, will very likely alter the conformation/dynamics of the system. Due to the larger systems and longer time spans allowed by CG MD, we chose this method to study if the above conclusions remain valid for a dodecamer and for 2 orders of magnitude longer simulation times. In particular, we calculated the free energy profiles of a dodecamer’s SU while it opens and closes. Consistent with the previous conclusions, we found that the differences in free energy are smaller for systems where the linker’s charge is partially or fully neutralized, thus rendering their transition to an open conformation more likely than for a wild-type SU.

The CG simulations also showed that the three systems studied, WT, PT286 and NE, have a tendency to increase the separation hub-kinase of some of their SUs. This was showed to increase the exposure of K21 and R451, residues that were found to interact strongly with Ca^2+^/CaM. Thus, a mechanism for CaM binding to T305/306 and opening of the system’s SU manifests:

- A SU (or several) of a CaMKII system increases its hub-kinase separation, partially exposing K21, but leaving T305/306 still mostly buried.
- If Ca^2+^/CaM is available, the now relatively exposed K21 is able to strongly interact with it, thus attracting it to its neighborhood, which holds T305/306.
- Finally, the strong interactions between CaM and the now near T305/306 (and nearby residues; see table in Figure 5) may result in their binding.

In summary, this study suggests a physical explanation for the mechanism guiding the activation of CaMKII’s subunits, in terms of electrostatic interactions. This is pictured in Figure 8, where the electrically charged patches are represented in red (negative) and blue (positive); Ca^2+^/CaM, phosphorylated residues and, possibly, cations in the solution, are the switches which turn on and off the interactions between patches. The activation process of CaMKII starts with a natural partial opening in the SUs (Figure 7) which, through exposure of positive residues K21 and R415, causes an attraction of Ca^2+^/CaM to them and the nearby T305/306. This may result in the binding of the latter two, and partial neutralization of the positively charged patch of the linker, thus favoring the SU opening (Figures 2 and left panel of 4). CaM interacts strongly with the linker, via the formation of hydrogen bonds (Figure 5), forbidding the re-closure due to its location between hub and kinase. This conformation, with the kinase far from the hub, increases the volume available for a neighboring kinase to get close to T286 and phosphorylate it. Later on, once CaM unbinds from the linker, the open kinase may associate to a neighboring hub (right panel of Figure 4, activation-competent conformations of Myers et al [38], also observed in [23]). Next, the small distance between the kinase and neighboring hub may be enough for a strong interaction leading to the cross-phosphorylation of T286 (and/or T305/306) of the hub’s SU. Finally, this phosphorylation partially neutralizes the charge of the patch and favors the partial opening of this neighboring SU, which exposes residues K21 and R145, initiating the activation process of this other SU.

**Figure 8:**
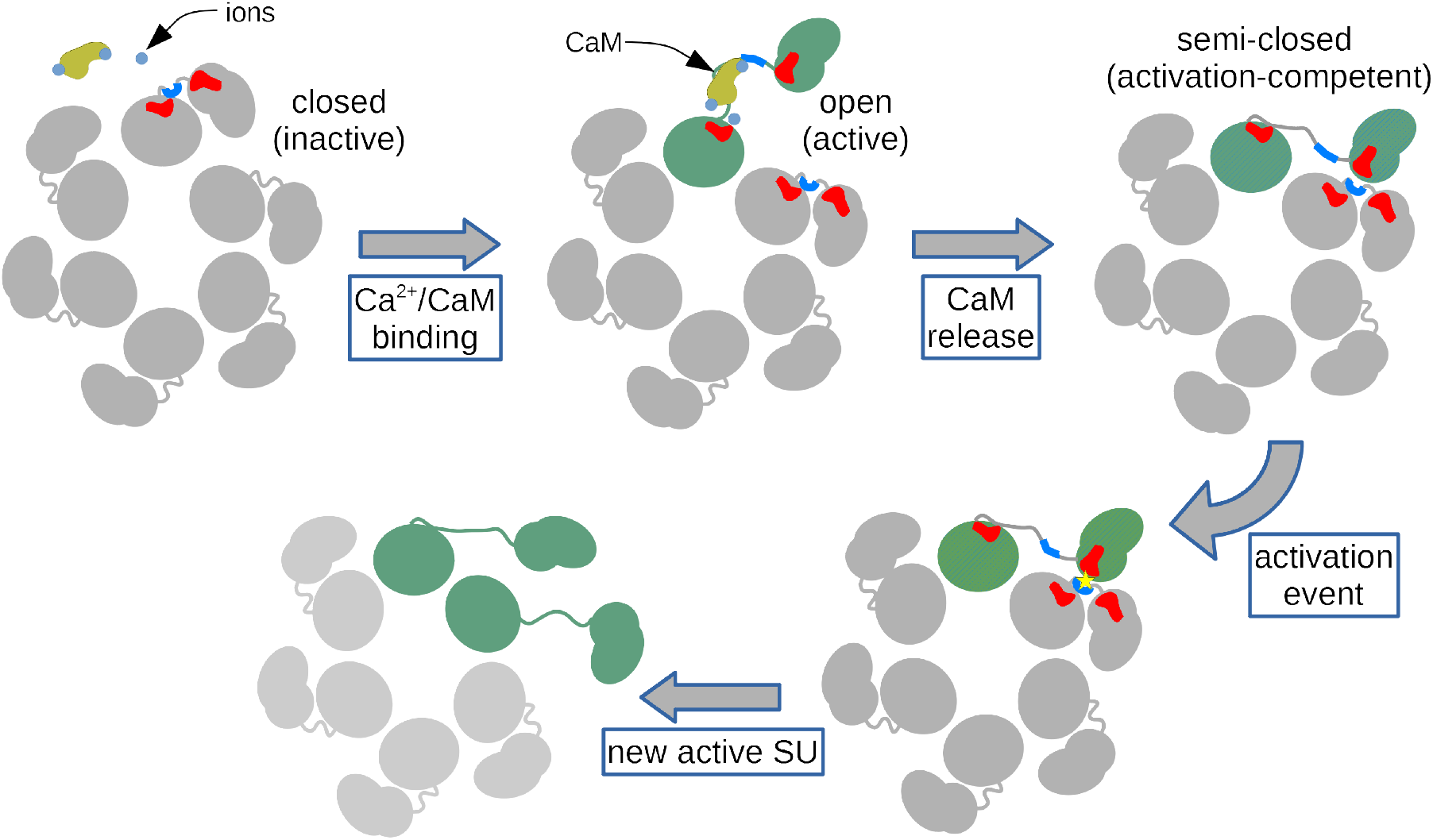
Steps in the activation of a CaMKII subunit. The first transition shows a closed (gray) SU (with negatively charged patches in red, and positively charged in blue) being affected by Ca^2+^/CaM (in yellow and light blue, respectively) and resulting in its opening (green SU). Once the complex is released, the SU is dynamically driven to a semi-closed conformation (green/white hatched SU), with its kinase domain getting close to a neighboring SU. The high negative charge of its kinase can interact with the high positive charge of the neighbor’s linker, thus favoring a strong interaction that could lead to the phosphorylation of the latter (activation event). This newly phosphorylated SU, now with a partially neutralized linker, is not favored as before to stay in a closed conformation (the previous attraction hub-linker-kinase was weakened). Thus, the systems acquires a new active SU.

The true hurdle to initiating the activation process is the opening of the first SU, which would be in a wild-type state and, thus, not exposing K21/R415 as much as in the non wild-type cases. But once this occurs, the opening of the other SUs should be more likely, since they would have a phosphorylated T286, and more exposed K21/R415. This explains how a cooperative mechanism of activation can arise [39, 38].

## 5. Conflicts of Interest

There are no conflicts of interest to declare

## 6. Acknowledgements

E.K.A. and I.J.G. acknowledge support from Agencia Nacional de Promoción Científica y Tecnológica, for grants PICT-2015-1706 and PICT-2015-3832, respectively. E.K.A. also acknowledges support from Agencia Nacional de Promoción Científica y Tecnológica, for grant number PICT-2016-4209. This work was partially funded by FOCEM (MERCOSUR Structural Convergence Fund), COF 03/11. S.P. belongs to the SNI program of ANII. I.J.G. also acknowledges help from Tristan Bereau with the use of the peptideB force-field [40] which, although not mentioned in the text, was the initial model with which coarse-grained systems were tested.

